# Ubiquitin Proteasome Pathway annotation in *Diaphorina citri* can reveal potential targets for RNAi based pest management

**DOI:** 10.1101/2021.10.11.464014

**Authors:** Will Tank, Teresa Shippy, Amanda Thate, Crissy Massimino, Prashant S. Hosmani, Mirella Flores-Gonzalez, Lukas A. Mueller, Wayne B. Hunter, Susan J. Brown, Tom D’Elia, Surya Saha

**Affiliations:** Division of Biology, Kansas State University, Manhattan, KS 66506, USA; Indian River State College, Fort Pierce, FL 34981, USA; Boyce Thompson Institute, Ithaca, NY 14853, USA; USDA-ARS, US Horticultural Research Laboratory, Fort Pierce, FL 34945, USA; Animal and Comparative Biomedical Sciences, University of Arizona, Tucson, AZ 85721, USA

## Abstract

Ubiquitination is an ATP-dependent process that targets proteins for degradation by the proteasome. In this study, we annotated 15 genes from the ubiquitin-proteasome pathway in the Asian citrus psyllid, *Diaphorina citri.* This psyllid vector has come to prominence in the last decade due to its role in the transmission of the devastating bacterial pathogen, *Candidatus* Liberibacter asiaticus (*C*Las). Infection of citrus crops by this pathogen causes Huanglongbing (HLB or citrus greening disease) and results in the eventual death of citrus trees. The identification and correct annotation of these genes in *D. citri* will be useful for functional genomic studies that aid in the development of RNAi-based management strategies aimed at reducing the spread of HLB. Investigating the effects of *C*Las infection on the expression of ubiquitin-proteasome pathway genes may provide new information regarding the role that these genes play in the acquisition and transmission of *C*Las by *D. citri.*

## Data Description

The ubiquitin-proteasome pathway (Figure 1) is responsible for the targeted degradation of most proteins in eukaryotic cells [1]. It regulates the concentration of proteins and degrades misfolded proteins. Ubiquitin modification is an ATP-dependent process. Ubiquitination is initiated by ubiquitin-activating enzymes (E1), which activate and transfer ubiquitin to ubiquitin-conjugating enzymes (E2).

**Figure 1.**
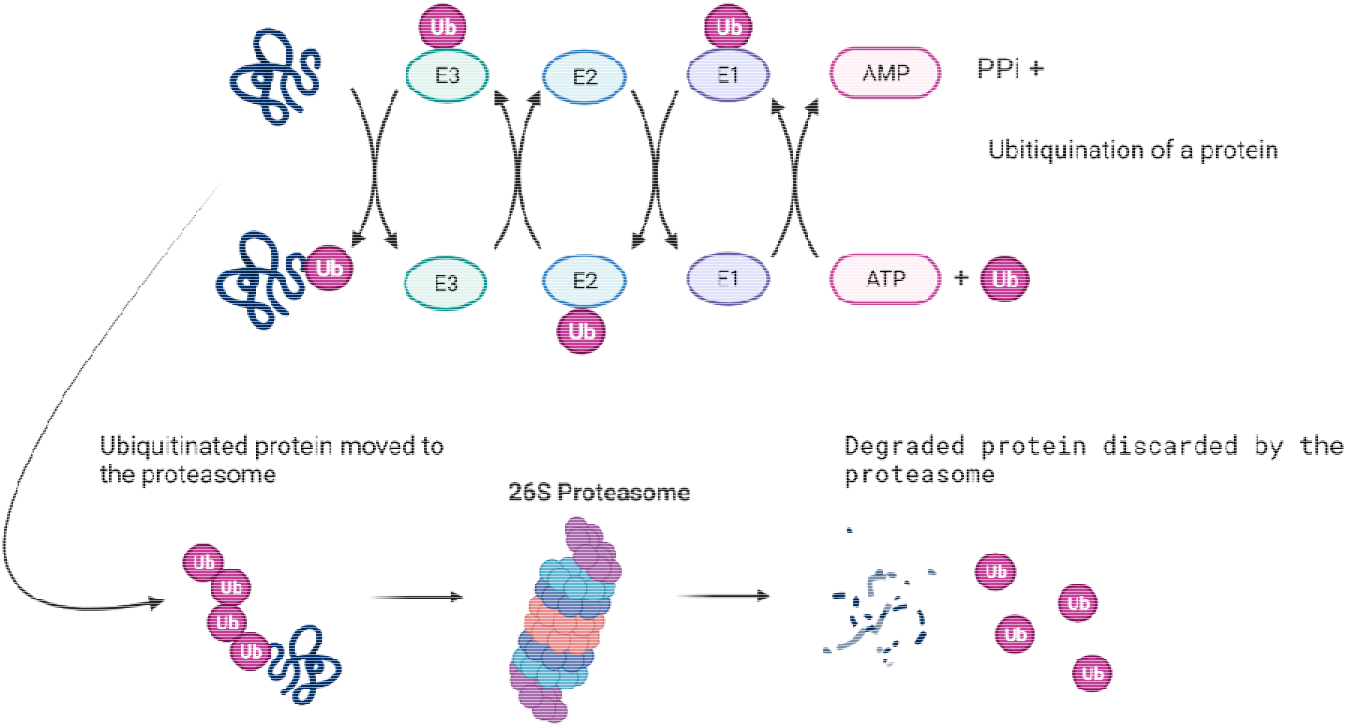
Ubiquitination process and interaction with the proteasome – Created with BioRender[3].

Conjugation of proteins with ubiquitin is mediated by ubiquitin ligases (E3) and tags the target protein for degradation by the proteasome. In its active form, the complete 26S proteasome consists of a 20S catalytic cylindrical core, and one or two 19S regulatory subunits bound to the termini on either or both sides of the core [2] (Figure 2).

**Figure 2.**
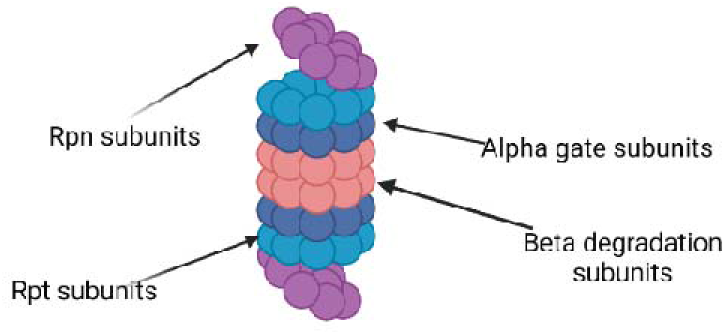
Proteasome and its subunits – Created with BioRender[3].

The 19S regulatory subunits, which are made up of Rpn and Rpt subunits, bind ubiquitinated proteins and may help unfold the proteins while translocating them to the 20S catalytic core. The 20S core, often referred to as the 20S proteasome, consists of two *β*-rings and two *α*-rings that organize into a cylinder-shaped quaternary structure. Proteolysis occurs at the N-termini of the *β*-rings where threonine residues form active sites that hydrolyze proteins via threonine nucleophilic hydrolysis [2].

## Context

The ubiquitin-proteasome pathway is highly conserved among eukaryotes and has been the target of many investigations attempting to decode the functions and inner workings of this pathway [1]. Studies involving arthropods [4] have linked the ubiquitin-proteasome pathway to development, immune response, gametogenesis, differentiation, apoptosis and stress response [5]. For example, the main pathways involved in insect resistance to viral and bacterial infections, Toll and JAK/STAT, are both dependent on proteasome activity for degradation of ubiquitinated targets [6,7]

In this study, we have identified and manually annotated 15 genes from the ubiquitin proteasome pathway in *Diaphorina citri,* which is the vector for *Candidatus* Liberibacter asiaticus (*C*Las), a bacterial pathogen associated with citrus greening disease. Several studies have suggested that the ubiquitin-proteasome pathway is affected by *C*Las infection of *D. citri.* Ramsey et al. [8] found that several ubiquitin-proteasome pathway proteins are differentially expressed between *C*Las(+) and *C*Las(−) *D. citri,* suggesting that protein degradation may be affected by *C*Las infection. Two other studies[9] reported differential expression of multiple E3 ubiquitin ligases during *C*Las infection [10]. With the purpose of finding possible pest control targets, Ulrich et al. [11] noted that a number of ubiquitin-proteasome genes were among the 40 genes associated with the highest lethality percentages in an RNAi screen of the beetle *Tribolium castaneum* [12], suggesting that this pathway is a possible target for RNAi-based control of insects.

## Methods

Ubiquitin proteasome genes in *D. citri* genome v3.0 [13] were identified through BLAST (NCBI BLAST, RRID:SCR_004870) searches of *D. citri* sequences with ubiquitin proteasome orthologs primarily from *Drosophila melanogaster.* Orthology was then verified by reciprocal BLAST of the National Center for Biotechnology Information (NCBI) [14] non-redundant protein database. Genes were manually annotated and validated in Apollo 2.1.0 (RRID:SCR_001936) using available evidence, including RNA-seq reads, Iso-seq, MCOT(MAKER (RRID:SCR_005309), Cufflinks (RRID:SCR_014597), Oases (RRID:SCR_011896), and Trinity (RRID:SCR_013048)) [13] (Table 2). A neighbor-joining phylogenetic tree using the CLUSTALW (ClustalW, RRID:SCR_017277) multiple sequence alignment with Poisson correction method and 1000 replicate bootstrap test was constructed in MEGA version 10 (Mega, RRID:SCR_000667) using full-length protein sequences from *D. citri, T. castaneum* [12] and *D. melanogaster* [7].

**Table 1.**
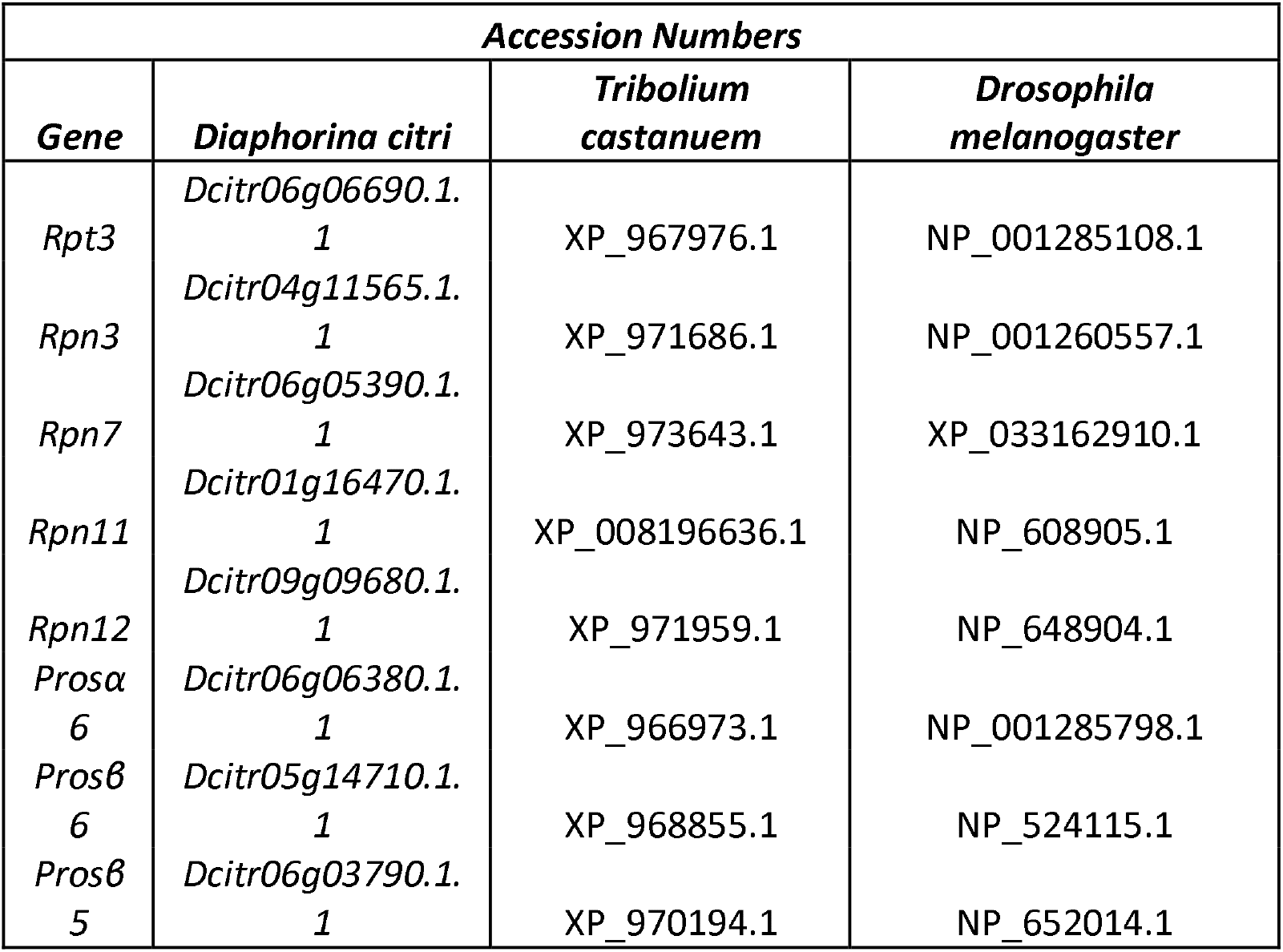
NCBI accession numbers for ortholog’s proteasome subunit annotations.

**Table 2.**
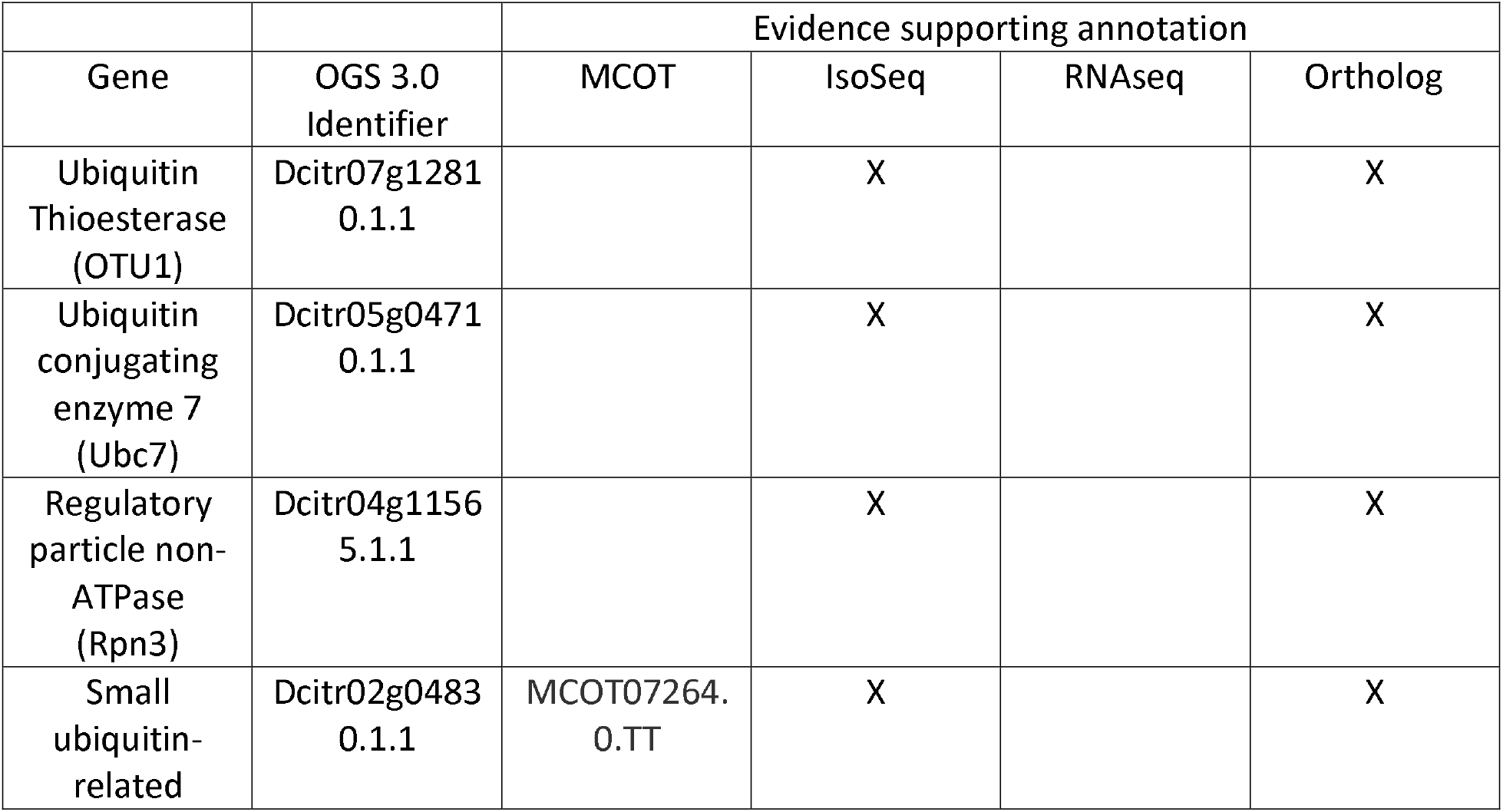
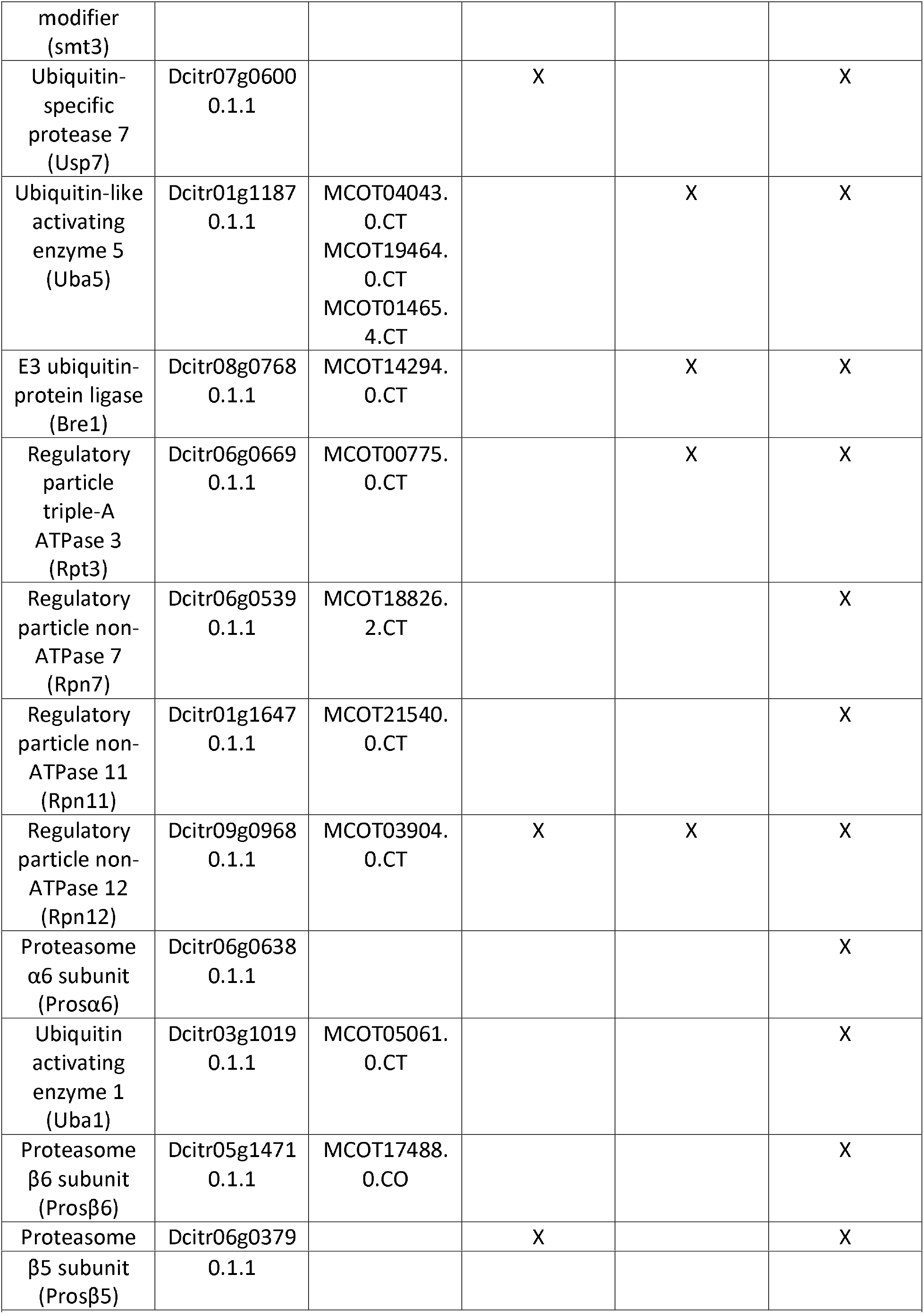
OGS v3.0 genes and the type of evidence used for validation of the gene model. Manually annotated ubiquitin-proteasome pathway genes in *Diaphorina citri.* Each gene has been assigned an OGS 3.0 ID and denoted as complete or partial. Evidence types supporting each annotation (MCOT, IsoSeq, RNAseq or ortholog) are indicated in the table. For the exact methods employed, see protcols.io document.

Expression data from the Citrus Greening Expression Network (CGEN) was visualized using the pheatmap [13] (pheatmap, RRID:SCR_016418) package of R [14] (R Project for Statistical Computing, RRID:SCR_001905) or Microsoft Excel (Microsoft Excel, RRID:SCR_016137).

Further details on the annotation process can be found at protocols.io [15] (Fig 3). Table 3 contains a list of orthologs evaluated for validation of the genes. Expression data from the Citrus Greening Expression Network (CGEN) was visualized using the pheatmap [16] (pheatmap, RRID:SCR_016418) package of R [17] (R Project for Statistical Computing, RRID:SCR_001905) or Microsoft Excel (Microsoft Excel, RRID:SCR_016137).

**Figure 3.**
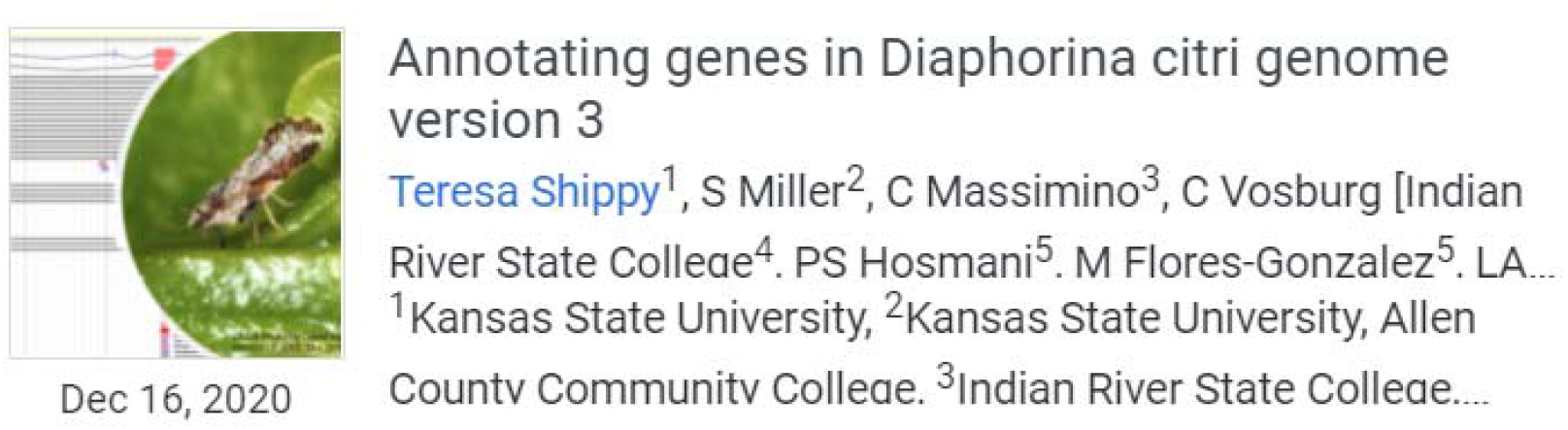
Protocols.io for psyllid genome curation[6].

**Figure 3.**
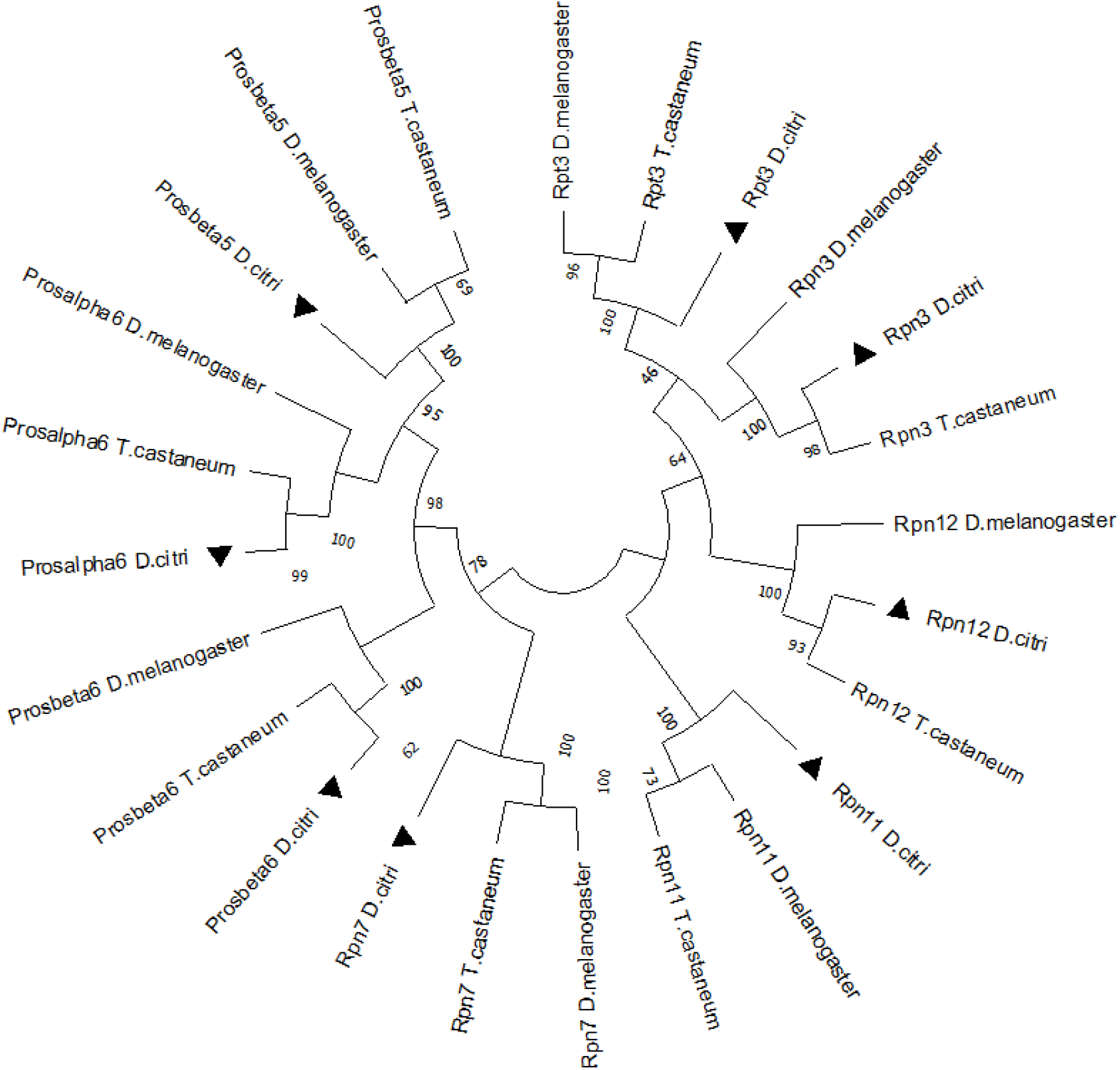
Phylogenetic comparison of orthologs

**Table 3:**
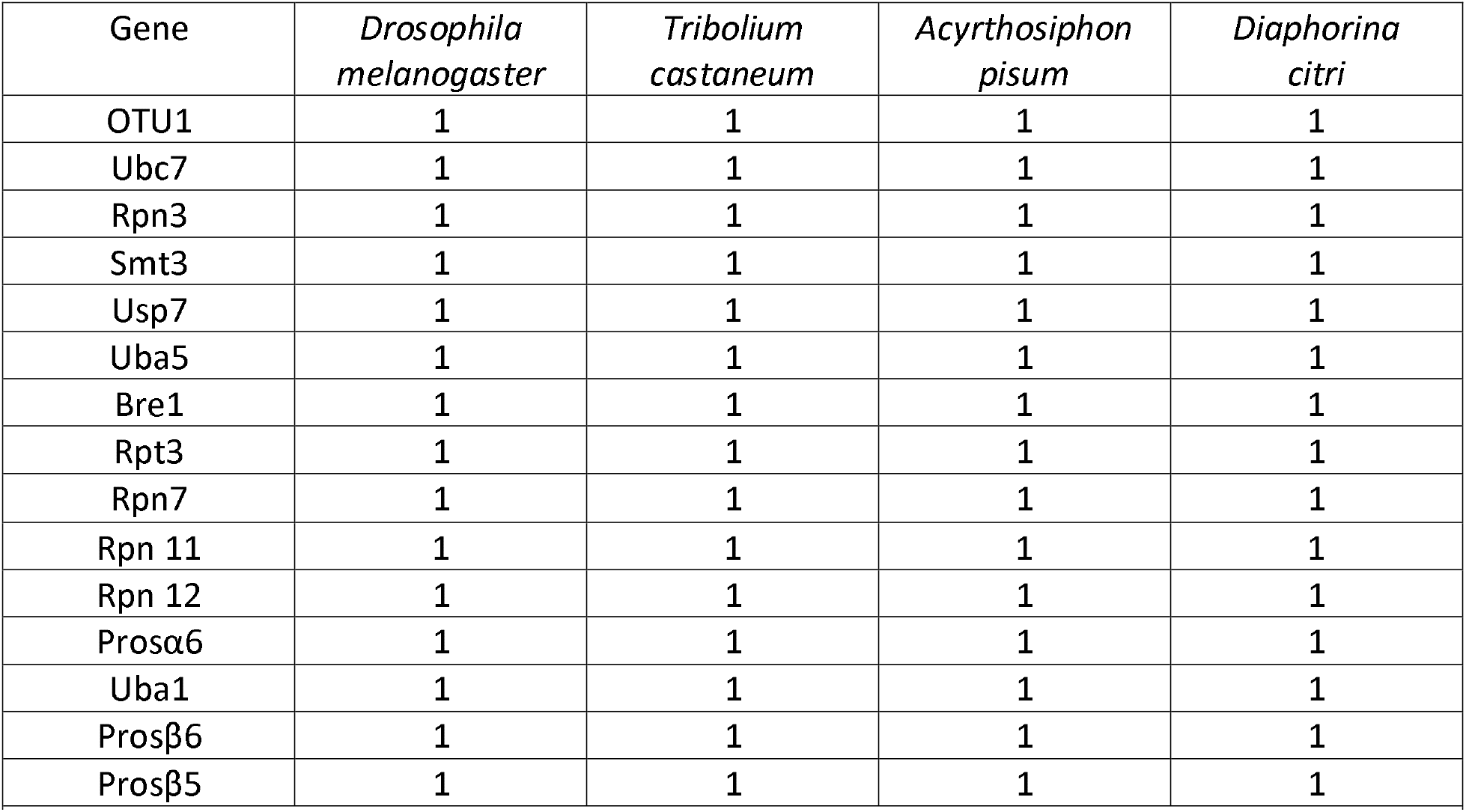
Count of orthologous genes in representative insects. *D. melanogaster* counts are from FlyBase [36] while *T. castaneum* and *A. pisum* counts were determined by BLAST searches.

### Data Validation and quality control

The ubiquitin proteasome pathway contains hundreds of genes. However, in our study we focused on the genes identified by Ramsey et al. [8] and Ulrich et al. [11]. We identified a single ortholog of each of the ubiquitin-proteasome pathway genes described in those studies (Tables 2 and 3), which is consistent with the gene count in *D. melanogaster* [7], *T. castaneum* [12] and *Acyrthosiphon pisum.* Genes were named based on their *Drosophila* orthologs. Since Ramsey et al. [8] observed differences in protein levels between *C*Las(+) and *C*Las(−) insects for the products of several of these genes, we used the Citrus Greening Expression Network (CGEN) [18] to compare transcriptional expression data from publicly available RNA-Seq samples of *C*Las(+) and *C*Las(−) *D. citri.*

Two main groups of genes were annotated in this study: ubiquitination-related genes and proteasome-related genes that encode proteasome-subunits.

### Ubiquitination-related genes

#### Ubiquitin activating enzyme 1 (Uba1)

As the only E1 enzyme in the ubiquitination pathway of insects, Uba1 is critical for initiation of the ubiquitination process [19]. In *D. melanogaster,* clones of cells homozygous for mutations in *Uba1* show changes in regulation of apoptosis, with weak alleles causing reduced apoptosis and strong alleles causing increased apoptosis [19]. Cells adjacent to clones for strong alleles of Uba1 overproliferate because of increased Notch signaling, suggesting that Uba1, through its role in ubiquitination, may act as a tumor suppressor gene. Flies completely lacking *Uba1* do not survive. However, flies homozygous for weaker alleles sometimes live to adulthood, but have reduced lifespan and motor defects[20]. Knockdown of *Uba1* in *Tribolium* also causes high levels of lethality [11], suggesting that *Uba1* could be a good target for RNAi-based pest control.

#### Ubiquitin conjugating enzyme 7 (Ubc7)

Ubc7 is one of about 30 E2 ubiquitin-conjugating enzymes known in *D. melanogaster.* It is also known as *courtless,* becauses male flies lacking Ubc7 display abnormal courtship behavior as well as defective spermatogenesis[21] In *D. citri,* Ramsey et al. [8] observed almost 6-fold higher expression of Ubc7 protein in *C*Las(+) organisms (Fig 4). To determine if this difference could be caused by changes in transcription, we analyzed *Ubc7* expression in publicly available RNA-Seq data in CGEN. Comparison of *Ubc7* transcript levels in *C*Las(+) versus *C*Las(−) samples did show slight upregulation of *Ubc7* in some **C*Las*(+) samples.

**Figure 4.**
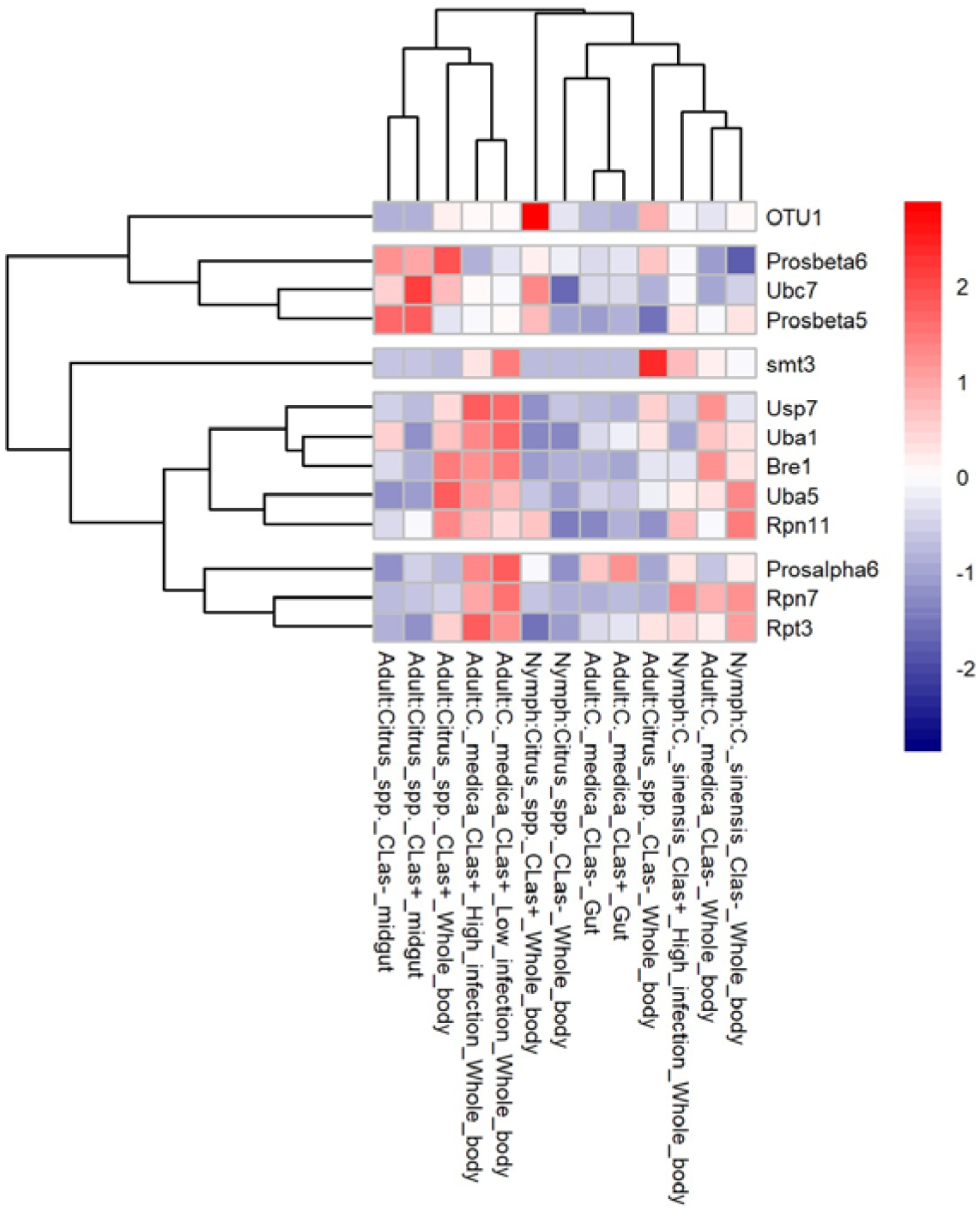
Differential expression heatmap scaled by gene.

#### E3 ubiquitin-protein ligase (Bre1)

As one of the E3 ligase genes, *Bre1* is a highly conserved gene affecting several other pathways through its role in the ubiquitin pathway. For example, in *D. melanogaster,* the Notch signaling pathway requires Bre1 for histone modification [22]. In *C*Las(+) psyllids, Bre1 protein levels were greatly reduced (almost 50-fold) compared to *C*Las(−) psyllids. However, no significant change in transcript levels is seen in the available RNA-Seq data (Fig 4)

#### Ubiquitin Thioesterase (OTU1)

In *D. melanogaster,* Ubiquitin Thioesterase 1 (OTU1) is a hydrolase responsible for removal of conjugated ubiquitin and regulates protein turnover by halting the ubiquitination process and retaining proteins [23]. In *D. citri,* Ramsey et al. [8] observed differential expression of the OTU1 protein during *C*Las infection with *C*Las(+) psyllids showing 13-fold higher expression than *C*Las(−). In nymphs fed on *C*Las(+) *Citrus* spp., we saw an approximately 6-fold increase in OTU1 transcripts compared to nymphs fed on uninfected plants of the same species. However, this was a single replicate study and none of the other *C*Las(+) versus *C*Las(−) comparisons showed a significant difference in OTU1 expression. (Fig 4).

#### Ubiquitin-specific protease 7 (Usp7)

Usp7 is another deubiquitinase/protease, and has been demonstrated in *D. melanogaster* to regulate aging and autophagy [24]. In *D. melanogaster,* specimens with knocked down Usp7 have difficulty coping with oxidative stress and starvation [24]. Furthermore, Usp7 has shown to interact with the p53 tumor suppressor gene [25] and inhibition of Usp7 through small molecule targeted approaches have shown promise in increasing cytotoxicity and inducing tumor cell death [25]. In contrast to OTU1, which also negatively regulates ubiquitin-mediated degradation, Usp7 protein showed 3-fold downregulation in *C*Las(+) organisms[8] We saw no evidence of differential expression at the transcript level (Fig 4).

#### Ubiquitin-like activating enzyme 5 (Uba5)

Uba5 is an E1 enzyme for the ufmylation pathway which modifies proteins in a manner similar to the ubiquitination pathway[26]. In ufmylation, the ubiquitin-like protein Ubiquitin fold modifier 1 (UFM1) is conjugated to target proteins. The functions of ufmylation are not well understood, but it seems to be important for tumor suppression, response to DNA damage, and regulation of protein homeostasis in the endoplasmic reticulum[27] Absence of Uba5 in *D. melanogaster* causes cerebellar ataxia, resulting in severe locomotive defects and a shortened lifespan [28]. Uba5 shows an almost 3-fold reduction in protein levels in *C*Las(+)compared to *C*Las(−) psyllids[8] However, there does not seem to be a consistent effect of *C*Las infection on *Uba5* transcript levels (Fig 4).

#### Small ubiquitin-related modifier (smt3)

The *smt3* gene encodes the Small Ubiquitin-Like Modifier (SUMO) protein [28]. Conjugation of SUMO to target proteins, known as sumoylation, occurs by a process analogous to ubiquitination but uses a distinct set of E1, E2 and E3 enzymes. Sumoylation has the opposite effect of ubiquitination in that it inhibits protein degradation [29,30]. Studies in *D. melanogaster* have shown that sumoylation may potentiate the immune response against infections [29]. Ramsey et al [8] observed a 4-fold reduction in *D. citri* Smt3 protein in *C*Las(+) versus *C*Las(−) psyllids. We do see a reduction in *smt3* transcript levels in adult psyllids fed on *Citrus* spp. infected with *C*Las, but this difference is not seen in nymphs or in adults fed on *Citrus medico* (Fig 4).

### Proteasome genes

#### Regulatory particle non-ATPase (Rpn3), Regulatory particle non-ATPase 7 (Rpn7), Regulatory particle non-ATPase 11 (Rpn11), Regulatory particle non-ATPase 12 (Rpn12)

The Regulatory particle non-ATPase (Rpn) subunits of the proteasome are responsible for the “lid” complex of the proteasome. They are directly responsible for capturing polyubiquitinated proteins and moving them into the cylindrical complex [2]. Furthermore, Rpn11 has been recognized as responsible for deubiquitinating these proteins. RNAi knockdown of Rpn subunits in *Nilaparvata lugens,* a member of the same insect order as *D. citri,* caused multiple defects in female reproduction, suggesting that the proteolytic activity of the proteasome is important for these processes [31]. Moreover, Rpn7, Rpn 11 and Rpn12 were shown by both Ulrich et al. [11] and Knorr et al. [32] to be potential high lethality RNAi targets in *T. castaneum.*

#### Regulatory particle triple-A ATPase 3 (Rpt3)

The six Regulatory particle triple-A ATPase (Rpt) subunits are organized into a hexameric ring, responsible for unfolding the proteins and opening the α gates within the proteasome, where the protein will be ultimately degraded [2]. Rpt3 is one of the three subunits which contains the conserved C-terminal hydrophobic-tyrosine-X (HbYX) [2], which has been shown to be directly responsible for facilitation of the gate opening. In fission yeast, Rpt3 has been shown to be associated with CENP-A, a variant of histone H3 responsible for kinetochore establishment [33,34]. In RNAi knockdown experiments in *Tribolium, Rpt3* was one of the top 11 genes with respect to lethality percentage.

#### Proteasome α6 subunit (Prosα6), Proteasome ß6 subunit (Prosß6), Proteasome ß5 subunit (Prosß5)

The Proteasome (Pros) alpha proteins are the “gating” subunits, which receive the unfolded proteins provided by the Rpt subunits [2]. The Pros beta subunits provide the peptidases and catalytic enzymatic activity that degrade the proteins within the proteasome. In *D. melanogaster,* mutations in the Pros units cause accentuated muscle degradation and a 25% decrease in lifespan [35]. The three Pros subunit genes annotated here were identified as good targets for RNA-based pest control because of high lethality percentages [11].

The protein sequences of the various proteasome subunits are often quite similar, making it challenging to establish orthology. We performed phylogenetic analysis to ensure that we had correctly identified all of the annotated *D. citri* proteasome subunit genes (Figure 3). All *D. citri* subunit proteins clustered with the expected *D. melanogaster* and *T. castaneum* orthologs, indicating that the annotated genes are named correctly.

In Ramsey et al [8], a very distinct difference can be observed between upregulated and downregulated proteins, with as much as 50 fold differences in expression. When these data are displayed as a clustered heatmap scaled by gene (Fig 4) we can compare the relative expression levels between samples. There is very little differential expression of genes between *C*Las(+) and *C*Las(−) samples. The apparent differences between *C*Las effect on protein and gene expression could indicate that there is post-transcriptional regulation of these genes/proteins. Post-transcriptional regulation of proteasome genes in yeast [25] has been observed before and could be a possible explanation for these discrepancies.

### Conclusion and future work

The ubiquitin proteasome pathway is highly conserved in most eukaryotes [1] and, therefore, a potential candidate for comprehensive studies. Disruption of this pathway has shown promise as a pest control mechanism in various insects, as explored by Ramsey et al [8], Ulrich et al [11], among many others, leading us to annotate 15 ubiquitin proteasome pathway genes from *D. citri.* Previous reports also suggested that the pathway might be affected by *C*Las infection[8,9]. Our analyses found little impact of *C*Las infection on transcription of the subset of ubiquitin-proteasome genes we annotated. More comprehensive studies will be needed to understand how *C*Las influences this pathway in *D. citri.*

### Re-use potential

The models annotated in this work will be helpful to researchers studying this pathway in many organisms, as this pathway is present in the vast majority of the eukaryotic species. Future studies are required to confirm the validity of the ubiquitin proteasome pathway as an RNAi target for control of *D. citri* populations.

### Data Availability

The *D. citri*genome assembly, official gene sets, and transcriptome data can be downloaded from the Citrus Greening website. Additional datasets supporting this article are available in the GigaScience GigaDB repository. [Ref to be added by editor]

## List of Abbreviations

*D. melanogaster*: Drosophila melanogaster;
*T. castaneum*: Tribolium castaneum;
*D. citri*: Diaphorina citri:
*C*Las: Candidatus Liberibacter asiaticus;
NCBI: National Center for Biotechnology Information,
CGEN: Citrus Greening Expression Network;
RNAi: RNA interference;
RNA-seq: RNA sequencing;
Iso-seq: Isoform sequencing;
MCOT: Maker, Cufflinks, Oasis, Trinity.

## Declarations

### Ethical Approval

Not applicable.

### Consent for publication

Not applicable.

### Author contributions

WBH, SJB, TD and LAM conceptualized the study; TD, SS, TDS and SJB supervised the study; SJB, TD, SS and LAM contributed to project administration; WT, AT and CM conducted the investigation; PH, MF-G and SS contributed to software development; PH, MF-G, SS, TDS and JB developed methodology; SJB, TD, WBH and LAM acquired funding; WT prepared and wrote the original draft; TD, SJB, SS, TDS, CM,WBH and JB reviewed and edited the draft.

